# An Algorithm to Build a *Multi-genome* Reference

**DOI:** 10.1101/2020.04.11.036871

**Authors:** Leily Rabbani, Jonas Müller, Detlef Weigel

## Abstract

1

**Motivation:** New DNA sequencing technologies have enabled the rapid analysis of many thousands of genomes from a single species. At the same time, the conventional approach of mapping sequencing reads against a single reference genome sequence is no longer adequate. However, even where multiple high-quality reference genomes are available, the problem remains how one would integrate results from pairwise analyses.

**Result:** To overcome the limits imposed by mapping sequence reads against a single reference genome, or serially mapping them against multiple reference genomes, we have developed the *MGR* method that allows simultaneous comparison against multiple high-quality reference genomes, in order to remove the bias that comes from using only a single-genome reference and to simplify downstream analyses. To this end, we present the *MGR* algorithm that creates a graph (*MGR* graph) as a *multi-genome* reference. To reduce the size and complexity of the *multi-genome* reference, highly similar orthologous^1^ and paralogous^2^ regions are collapsed while more substantial differences are retained. To evaluate the performance of our model, we have developed a genome compression tool, which can be used to estimate the amount of shared information between genomes.

**Availability:** https://github.com/LeilyR/Multi-genome-Reference.git

## 2 Introduction

The number of fully re-sequenced genomes has increased exponentially during the last decade. It provides scientists with vast amounts of diverse data which presents a major challenge when attempting to completely extract information from it. Advanced sequencing technologies not only resulted in vast amounts of sequenced genomes but also increased the quality of them substantially. Previously, it might have been optimal to use a high quality genome as a reference, however as the quality of re-sequenced genomes has improved, this no longer makes sense. In addition, as has already been shown by *Schneeberger et al.*,^32^ even when additional genomes are not complete, integrating known variants in a graph helps to call variants from reads. Recently, several alternative approaches have been proposed^10, 18, 22, 23, 32^ in which variations are added to the reference to break the linearity of the reference, creating a graph structure.

Graphs have been widely used in the history of genome analysis from assembling^35, 37^ to aligning^18, 26, 31, 39^ and representing variations.^4, 22, 23^ A common type of graph that has been used for genome assembly and analysis is the *de Bruijn* graph.^11, 18, 20, 37^ A *de Bruijn* graph is a directed graph where nodes are substrings of length *k* (*k* − *mers*) and edges represent an overlap of length *k* − 1 between two adjacent nodes. Graphs that are produced with this approach are dependent of the value of *k*. Previous methods have used *de Bruijn* graphs as a reference for mapping sequencing reads^18, 20^ or for assembling genomes.^11, 37^

Another commonly used graph is the sequence graph.^22, 23^ *Nguyen et al.*^22^ have proposed a bidirectional sequence graph that can be used as a pangenome reference for a population of genomes. To construct it, a parametric method is applied to a set of aligned genomes to obtain an orienting and ordering of homologous blocks of sequences that maximally agrees with the input genomes. More recently, *Novak et al* ^23^ used a sequence graph to construct a collapsed representation of a set of genomes, which was then used as a reference to query sub-haplotypes^3^.

In other studies,^4, 19, 28^ the main focus has been on introducing variant sites in a graph in order to determine which variants test genomes carry. Others^24, 32^ have focused on proving the ability of a graph in capturing new variations as an alternative to a linear reference genome. *Novak et al.*^24^ have recently shown that adding the variants to the reference may improve the estimation of an individual genome.

A further challenge as a consequence of a large number of sequencing and re-sequencing research is finding a way to store the large amount of data that is produced or used utilising the new sequencing technologies. As DNA sequences contain only four different characters, effective compression should achieve an information content below 2 bit per base. However, standard text compression algorithms like gzip^3^ fail to achieve this on real biological sequences. Due to sequence duplication events, biological DNA sequences contain numerous similar regions. Standard text compression tools focus more on detecting exact repetitions and do not have adequate models for error-prone biological duplication processes. Several studies have been done to build a DNA specific compression tool. In some studies one or multiple reference genomes are used to compress the genome data,^2, 7, 30, 36, 38^ other tools work *de novo* where encoding and decoding are independent of a reference sequence.^9, 33^ Reference-based compressions return a higher rate of compression since they include prior knowledge about data. However, they have a drawback in that the same reference is always needed for decoding. Some studies focused on compressing short sequencing reads.^12, 33^ Whereas, with others such as in *iDoComp*^25^ the algorithm is designed to compress a full length genome. In fact in the latter, a reference-based method has been used and about 60% compression was reported in most of the cases.

Here, we introduce the *MGR* algorithm, a global approach to create a string graph as is defined by *E. W. Myers* ^21^ where the vertices are fragments of sequences while the edges present the order of the vertices on the genome. Our approach generates such a graph which can be used as a reference and represents several input genomes. As opposed to the above algorithms, we compress several full length genomes instead of a single reference genome and its variants. Shannon information^34^ is used as the cost function which makes the algorithm independent of the choice of any arbitrary parameter value. Higher order Markov models are used to estimate the information content of DNA sequences as well as of pairwise sequence alignments. The algorithm clusters sequences based on the information content of the sequences and alignments between them and selects a representative for each cluster. This clustering based approach allows for non-parametrically identifying important variation which will result in different cluster centres. Non-important variation will be encoded as differences between the representative sequence of a cluster and its members. The created *MGR* graph represents the data reliably and gives us the opportunity of traversing over it and finding a mosaic of genomes that fits a new sequence fragment the best. To evaluate our approach, we also propose a DNA compression tool that can be used to compress full length genomes by exploiting the long repetitive elements that are not identical. Thus, it can be an excellent tool to study differences between assembled genomes on a global scale.

## 3 Methods

The algorithm was implemented in C++.

### 3.1 Preparing the input data

The main goal of the algorithm is to find representative elements from each group of similar sequences (orthologous or paralogous regions) by minimizing Shannon information. Therefore, for the first step, similar regions within and between sequences need to be found. To this end, a single FASTA^17^ file is generated containing all the input genome sequences with a unique accession ID assigned to each input genome. This file will then be used as an input for a pairwise aligner tool to capture similar regions in the form of local pairwise alignments within and between input genomes. The alignment file in the Multiple Alignment Format (MAF)^1^ along with the generated FASTA file are provided to the algorithm as input for generating a *MGR* graph. Figure 1 shows an overview of the *MGR* algorithm operating on a given set of pairwise alignments.

**Figure 1:**
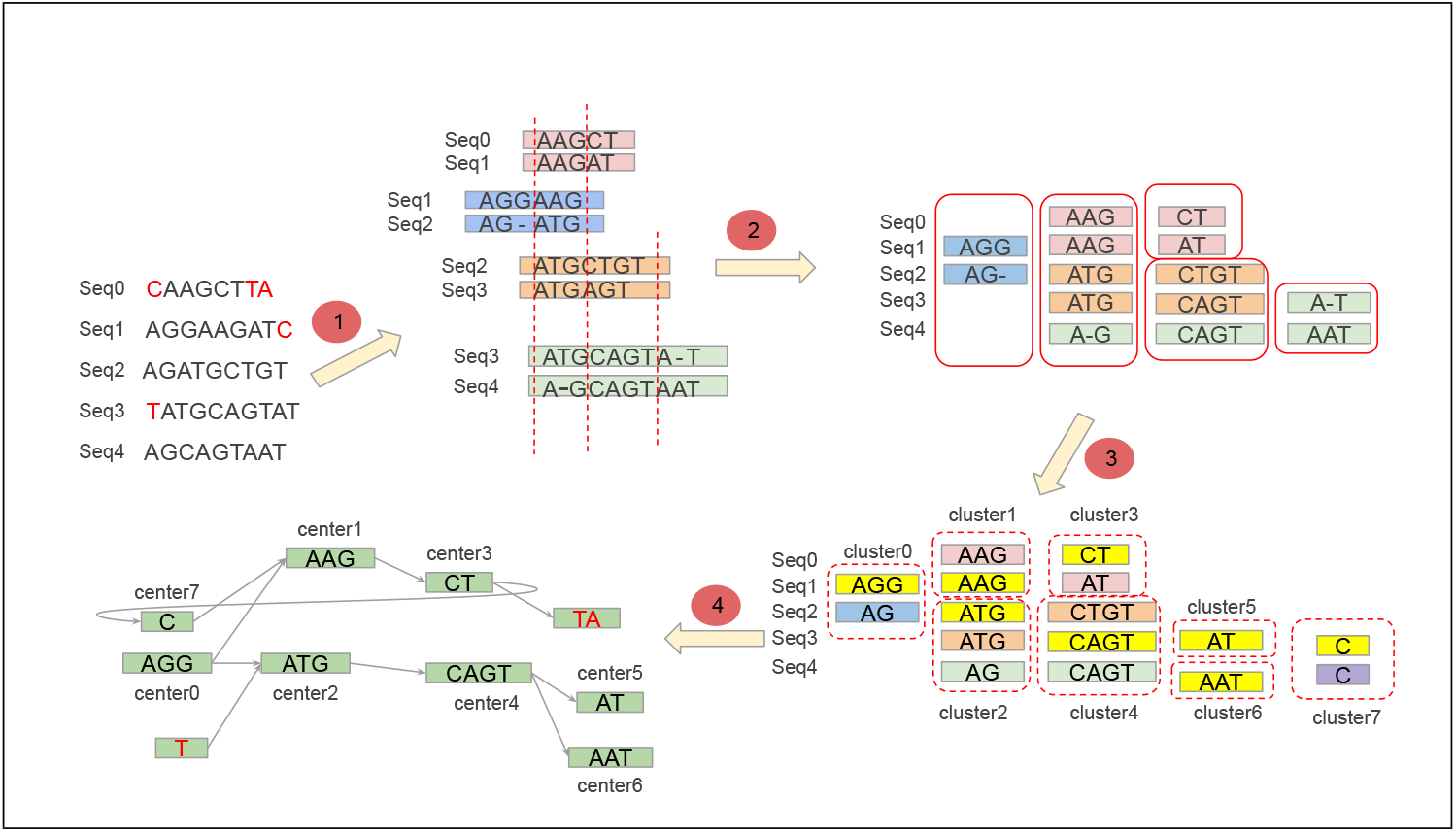
General overview of the *MGR* algorithm Step one: Detection of pairwise alignments. Step two: Division of alignments into non-overlapped segments. Step three: Grouping of related sequences into different clusters. Step four: Converting cluster centres into vertices of the graph.

### 3.2 Training the model

We use variable order Markov Chain (MC) models to compute the information content of sequences and pairwise alignments. In a first order Markov Chain model, the probability distribution of the next character depends solely on its predecessor. Such a model can be used to estimate the information cost of a sequence. If the probability distribution of the model is a good representation of pairwise character successions in the sequence, it will yield a low information cost. By making the probability distribution of the next character depend on a longer context, we create a higher order Markov Chain model. This can lower the information cost of a sequence if the context specific probability distributions represent the data well. Long stretches of approximate repeats, like in biological DNA sequences, can benefit from using higher order Markov Chain models. However, incrementing the order increases information content of a Markov Chain model exponentially. This creates a trade-off where longer contexts need the encoding of a high number of context specific probability distributions which lowers the information content necessary to encode the sequence. We solve this by using a greedy approach. Starting from contexts of length one, we look at the total cost to encode using a first order Markov Chain model and then the sequences using this model. Then we iteratively check whether increasing the order of the Markov Chain or just the length of a specific context lowers the total information cost. We also optimize the number of bits used to encode each individual probability distribution of the model contexts.

A sequence alignment can be interpreted as a series of instructions needed to modify one reference sequence of the alignment to the other one. These series of modification instructions can also be modelled using variable order Markov Chain models.

The probability distributions given by these models yield a lower bound of information cost necessary to encode each character or modification instruction. This is called Shannon information. It is the negative binary logarithm of a probabilistic event. Thus, an optimal encoding algorithm will require many bits to encode a rare event but only very few bits to encode a common event. Transforming probabilistic models to estimations of encoding efficiency makes sequence and alignments comparable. The information cost of both data types is measured using the same unit, the bit. This yields a non-parametric approach to decide whether an alignment is sufficiently important, or not. We check whether including the alignment into an encoding of all the sequences would reduce the total amount of Shannon information. Instead of just encoding all sequences using the sequence based variable order Markov chain models, we can incorporate a pairwise alignment. Thus, we only need to encode one of the reference sequences of the alignment. The other one can be defined by applying the modification instructions contained in the alignment to the first one (see Figure 2 for the illustration). If this approach yields a reduction of the total information content, we include the alignment into the model.

**Figure 2:**
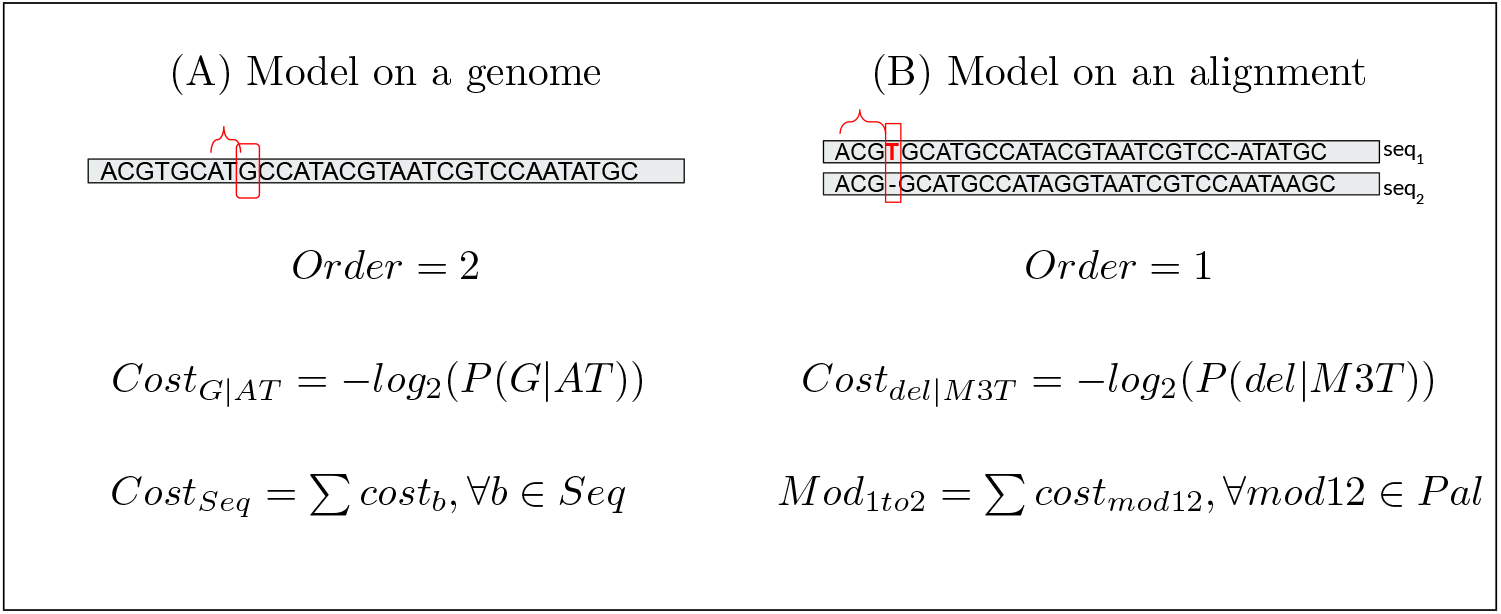
Simple illustration of how the models are trained. Red squares present the current position, where the model is trained. The highlighted base in (B) presents the current base, where the modification between two sequences of the pairwise alignment is read.

For this decision to be made, the gain values are computed for each alignment. These indicate how much information can be gained by using the alignment instead of two reference sequences separately. Formula 1 shows this calculation where *G*_1_ is the gain of creating the reference 2 of an alignment and modifying it to the reference 1, instead of creating both references. The opposite of the latter is the case for *G*_2_.

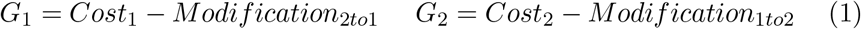

In a real encoding we cannot realize the complete amount of the information gain computed above. To ensure lossless compression, we also need to encode the following instructions: 1) Stop sequence based decoding and start alignment based encoding. 2) Index of the reference sequence on which the alignment will be based (we do not need a position given in base pairs as the actual implementation only uses cluster centres as other reference). 3) Stop alignment based decoding and switch back to sequence based decoding. We estimate the information cost of these encodings based on their expected frequencies and call their sum a base cost. An alignment is kept if its average gain is higher than a base cost; otherwise it is discarded.

### 3.3 Filtering homology

So far, potentially similar regions on different genomes are detected. Once these regions are successfully found, all the alignments are then reweighed. Weighing is done by considering how many times an alignment is confirmed by a third reference genome (Figure 3). The number of such references is counted for each alignment and the gain values are recalculated using this number.

**Figure 3:**
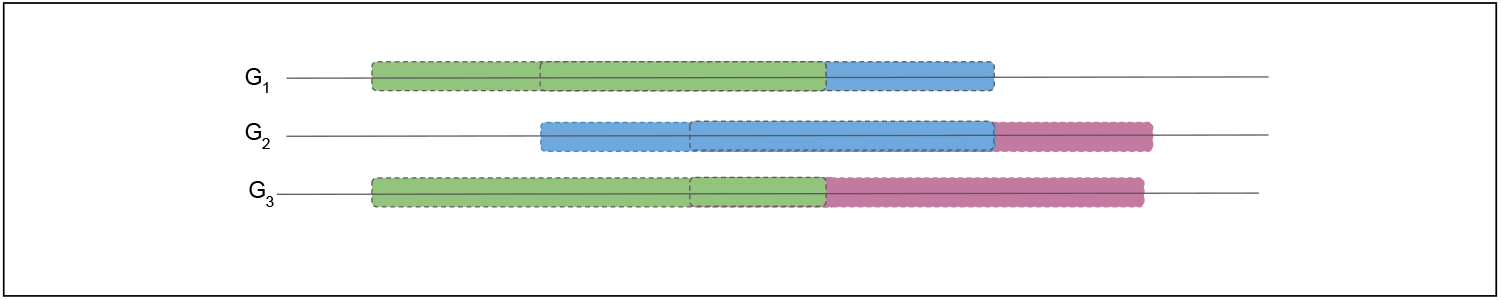
Searching for sequence similarities *G*_1_, *G*_2_ and *G*_3_ are three input genomes. The blue alignment is highly overlapped with the pink and the green alignments. That these regions are typically contiguous in a genome is confirmed by the coloured region on *G*_3_.

At the end only a fraction of the alignments, those with the highest gain value, are kept. The filtering can be considered as an optional step to reduce the input size by retaining more valuable alignments, which are the ones that have higher potential of containing a representative piece of sequence.

### 3.4 Removing redundant data

After filtering the data, one needs to deal with the partially overlapped alignments. They need to be detected and the redundancy has to be discarded otherwise one piece of a sequence might be taken into account more than once. In order to solve this conundrum, alignments are cut into smaller non-overlapped segments.

Redundant regions are propagated over a set of alignments and this makes the cutting process computationally expensive and time consuming. To handle this, cutting is done iteratively on different levels of redundancy. In each iteration, a graph of the overlapped alignments is generated and its 2–*edge* connected sub-graph is extracted. The alignments on this sub-graph are then cut into non-overlapped segments and saved in a data container along with the rest of the alignments on the generated alignment graph. After several iterations over different levels of redundancy the entire remaining alignments are checked for any level of overlap and their redundancy is discarded. To this end, alignments are grouped together if there is an overlap between them. Therefore, there will no redundancy between different groups of alignments but within them. At the end, each group of alignments is cut into non-overlapped segments of sequences.

Since the partial overlap has been eliminated, pairwise alignments can be assigned to different groups considering the complete overlap between one of their references. Sequence segments of each group are related to each other and are totally independent from the segments in the other groups.

### 3.5 Clustering sequences

Next, we cluster the sequences of each group in a way which minimizes the information cost. The AFP clustering algorithm^6^ is run over each group of independent pieces and assigns each sequence to a cluster.

One advantage of the AFP clustering algorithm is that it detects representatives of each cluster as well as the associated members, the other advantage of this algorithm over some other clustering algorithms which are commonly used in literature is that there is no need to set the number of clusters. It optimizes the total information cost to create all representatives as well as the information cost of alignment derived modification instructions to create cluster members from their representatives.

For each input genome sequence a series of all the centres and non-aligned regions that are not in any cluster is created. At the end, if there are non-aligned segments with the exact same content, within or between different input genomes, they will be grouped together and form a new cluster.

### 3.6 Building a *multi-genome* reference

A directed string graph as is proposed by *Myers*^21^ is created as an output of the program. The output graph *G* = (*V*, *E*) is constructed over a set of vertices *V* which represent the (not always exactly) repeated pieces on different genomes. Two vertices *v*1, *v*2 ∈ *V* are connected with an edge *e* ∈ *E*, if and only if they are next to one another on at least one of the input genomes (Figure 1). This graph can be used as a *multi-genome* reference to remove the bias against a single reference.

The output graph is saved as a DOT^5^ and a GFA^15^ file format that shows all the adjacent vertices. The content of all the centres are also saved as a FASTA^17^ file, along with a text file that contains the extra information about the origin of each centre. Moreover, each cluster is saved on a MAF^1^ file as a multi sequence alignment between the centre and all its members.

### 3.7 Data compression

The constructed graph is then used to compress all the input genome sequences. During training, the model parameters, including the cost of creating each letter on a sequence as well as the cost of each modification pattern of an alignment are written on a file. After clustering, the indices of all the centres are added to the same file and the centres’ content are arithmetically encoded.^14^ In this step, using the clustering result, all the sequences are arithmetically encoded and the encoded result is written on the same file. For this purpose, each base on a sequence is encoded as long as there is no alignment on that position of a sequence. As soon as an alignment is reached, the index of its corresponding centre and the modification instructions for modifying each base of the centre to the corresponding base on the input sequence are encoded. The higher the compression rate the better the model fits the input data. Thus, it can be used as a measurement to evaluate the trained models.

## 4 Result

Several sets of data with different numbers of input genomes of various lengths were applied to assess the performance of the algorithm. In each case, a single FASTA file has been generated by the *MGR* algorithm containing the unique assigned accession ID to each input genome which makes the FASTA files compatible to what the algorithm needs for the next steps. For our experiments, we have always run LAST^13^ as an aligner on each of the generated FASTA files to create local pairwise sequence alignments, however, the *MGR* algorithm is not dependent on the choice of the aligner. The local pairwise alignments between and within sequences detected by this program were saved as a MAF file. Each of the MAF files and the their corresponding generated FASTA files have been used as input for the *MGR* algorithm. The size of the built *MGR* graph and the efficiency of the models, based on the obtained compression rates, were examined to measure the performance of the algorithm.

The pilot experiment was done on several strains of *E. coli* with the length of almost 5*Mb* per genome (Data section). Input files of different sizes were generated to evaluate the time complexity of the method. Four FASTA files with two, four, eight and sixteen input genomes of *E. coli* were built. After creating the corresponding MAF files for each of them, the number of alignments as well as their average length have been computed (Table 1). Relationships between the number of alignments normalized by their average length, and the run time of the program is then used to estimate the complexity of the algorithm (see Figure 4).

**Table 1:**
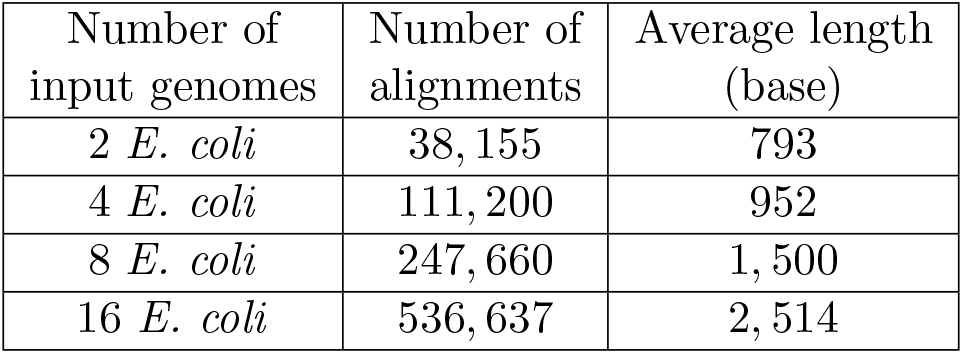
The number and the average length of the pairwise alignments on 4 different input dataset.

**Figure 4:**
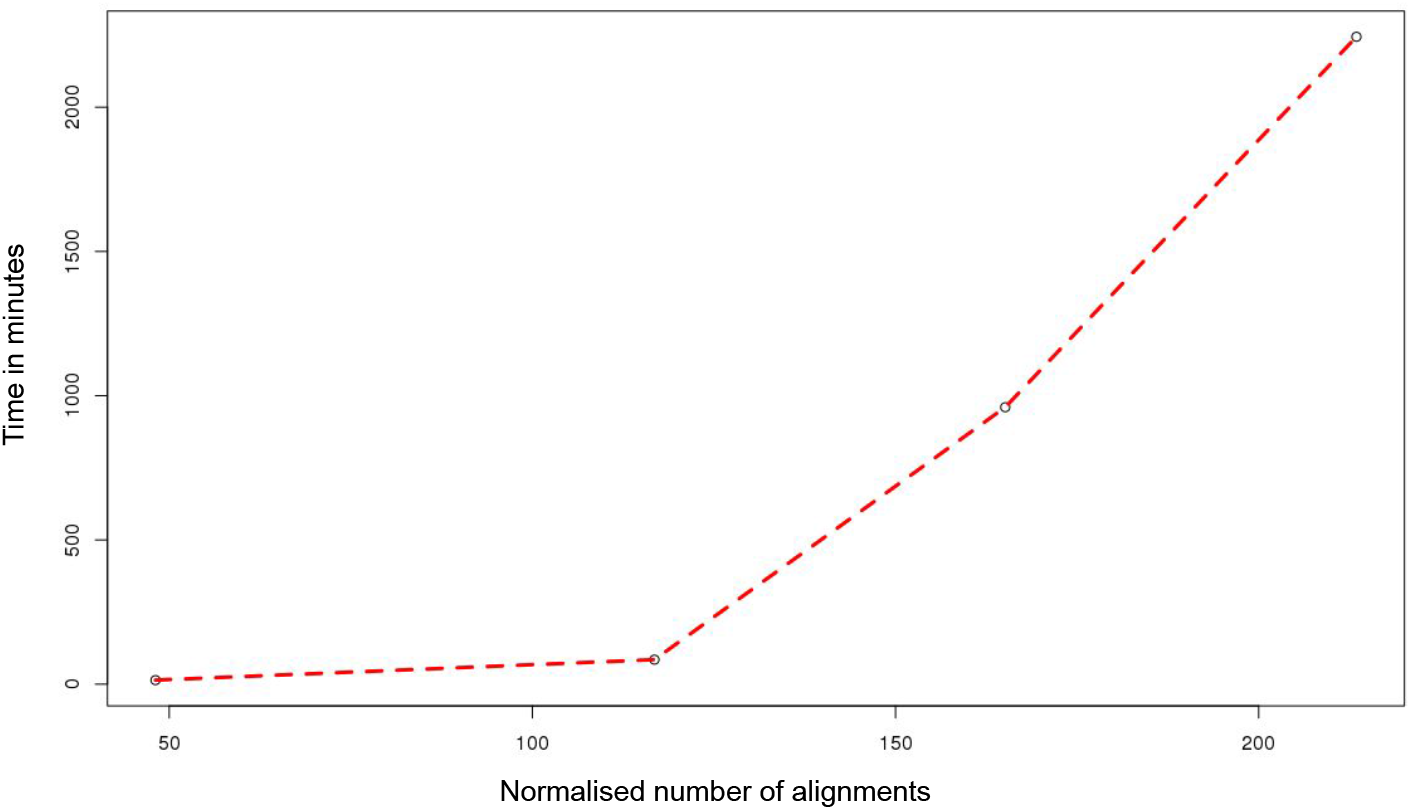
Relation between the number of input alignments normalised by their length and the running time.

On the same set of data, the compression rate computed by the *MGR* algorithm is compared with the compression rate obtained from a known genome specific compression tool, *MFcompress*.^29^ Close rates to what has been achieved by *MFcompress* (see Table 2) reveals the efficiency of the *MGR* algorithm in capturing the repetitive regions.

**Table 2:**
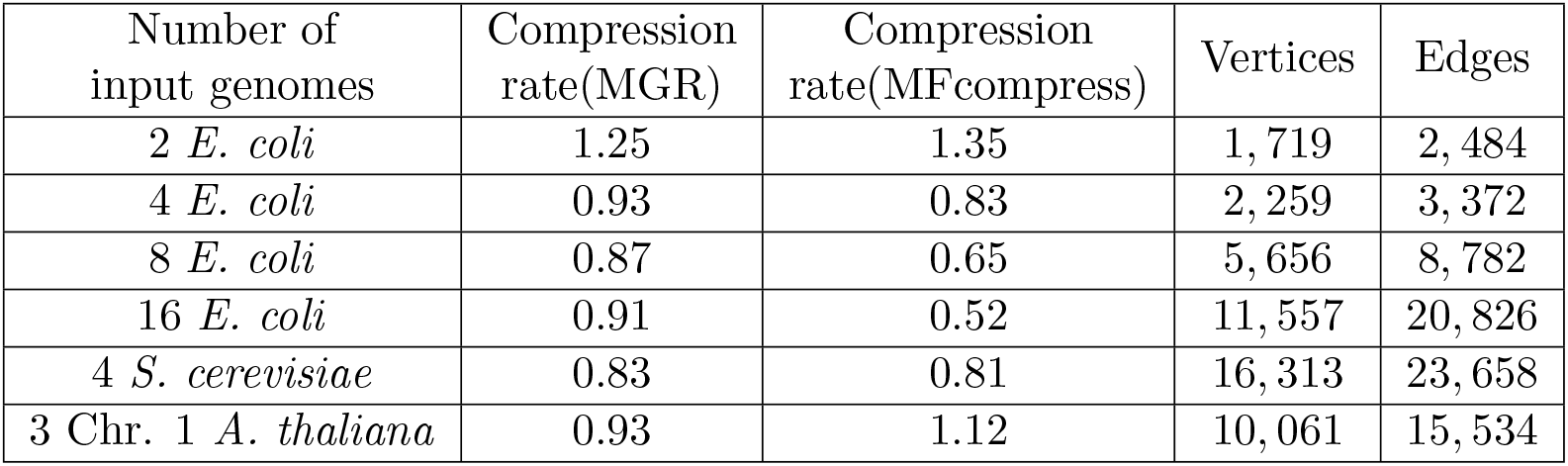
Comparison between the compression rate obtained from our algorithm and *MFcompress* as well as the number of vertices and edges on the generated *Multi-genome reference* graph

To ascertain the effectiveness of the program in processing larger genomes, the next experiments were carried out on larger input files. The first contained the genome of eight *Saccharomyces cerevisiae* strains with the length of 12*Mb* per strain while the second contained chromosome one from three different *A. thaliana* strains with the sequence length of 30*Mb* per chromosome (see Data section). The same procedure that has been used for the pilot experiment has been applied on each of the above mentioned datasets to generate the compatible input files and to run the algorithm on them. The size of the built graph in each case as well as the compression rate obtained for each of them are shown in Table 2. In all the cases, the high compression rate achieved reveals that the trained models fit the data well and that capturing more repetitive regions – as in the more complex S. cerevisiae genomes – raises the compression rate.

Moreover, to prove the simplicity of the data structure that the *MGR* algorithm offers in presenting several genomes, the size of the built *MGR* graph on two *E. coli* genomes has been compared with the size of created graphs by the vg tool^8^ and the REVEAL program^16^ on the same set of data. Table 3 shows this comparison where the number of vertices and edges are significantly lower on the *MGR* graph.

**Table 3:** Size of the graphs created by our tool, the vg tool and the REVEAL program on two *E. coli* genomes.

| Input<br>genomes | Vertices<br>(MGR) | Edges<br>(MGR) | Vertices<br>(Vg) | Edges<br>(Vg) | Vertices<br>(REVEAL) | Edges<br>(REVEAL) |
| --- | --- | --- | --- | --- | --- | --- |
| 2 *E. coli* | 1,719 | 2,484 | 415,701 | 374,468 | 91,296 | 121,798 |

## 5 Discussion

As previously stated, advanced sequencing technologies improved the quality as well as the quantity of sequenced genomes drastically. In conjunction with this, it has already been shown^24, 27, 32^ that adding extra information to a single genome results in a higher quality of mapping and detection of variations. Thus, shifting from a single reference genome to the *MGR* graph as a reference not only offers a robust data structure to present several genomes but also offers a reliable method to call the variants and to decrease the bias against using only a single reference.

Moreover, developing such an algorithm could assist us in searching for longer sequencing reads and for finding the best match for each read by walking through the graph and locating the path that fits the sequencing read the best. In contrast with the Schneeberger et al.^32^ approach, the *MGR* algorithm does not return a single genome as the closest reference for the mapped sequencing reads but it is incapable of finding a new genome which is closer to a mixture of several existing genomes.

To our knowledge the *MGR* algorithm is one of the first algorithms which takes advantage of several fully sequenced genomes as opposed to using variations along with a single reference genome. Given a piece of sequence, the *MGR* algorithm estimates its information content *de novo* or estimates it from an already known piece of a sequence. Estimating from a known piece can be done by looking for the information content of the changes between the given sequence and the known piece of sequence. The higher order Markov models used in the *MGR* algorithm estimate the information content of DNA sequences as well as of pairwise sequence alignments reliably. Since decisions are always made in a manner to minimize the Shannon information^34^ the algorithm does not rely on arbitrary parameter choices. As well as the above mentioned criteria, applying genome specific Markov chain models allows us to analyze genomes with very different magnitudes of difference together.

Another advantage of applying the *MGR* algorithm compared to the other methods is that it clusters sequence pieces. Given a set of sequences, between which pairwise alignments exist, *MGR* selects some of them as representatives (with *de novo* information cost). For this, the Affinity Propagation Clustering (AFP) algorithm^6^ is used, which automatically approximates the optimal number of clusters by minimizing the total cost needed to create all input elements either *de novo* or based on one of the representatives. Although clustering results in the discarding of unimportant variation by collapsing highly similar repeats, all the variation can still be accessed if necessary. Moreover, it enables us to perform both between and within cluster analysis and can be easily applied in global genome analysis and represents their diversity. It is especially helpful in studying repeats. Using clustering also resulted in a simpler representation of several genomes compared to existing methods such as REVEAL^16^ and vg tool.^8^

The applied DNA compression in the *MGR* uses the same strategy of minimizing Shannon information^34^ to actually compress the input sequences and does not rely on using similarities between each DNA sequence and a certain reference genome. The compression performance of the algorithm not only shows how efficiently the Markov chain models capture the intrinsic properties of DNA sequences but can also be used as a metric to measure the common information of individual genomes and shows their distances. Thus, it can be an excellent tool to study differences between assembled genomes on a global scale.

## Data

Data sets which were used to build a *multi-genome reference* over *E. coli* genomes are the following 16 NCBI reference sequences.

NC 000913.3, NC 002695.1, NC 010473.1, NC 012967.1, AE014075.1, NC 004431.1, NC 007946.1, NC 008253.1, NC 011750.1, NC 011751.1, NC 018658.1, NC 009800.1, NC 017634.1, AE005174.2, NC 008563.1, NC 017633.1

The *S.cerevisiae* strains can be found here:

https://www.yeastgenome.org/strain/AWRI1631
https://www.yeastgenome.org/strain/FL100
https://www.yeastgenome.org/strain/CLIB215
https://www.ncbi.nlm.nih.gov/Traces/wgs/?val=AEWL01#contigs

*A. thaliana* Data:

PRJEB37261 Ler-0 (AT7213)
PRJEB37258 Ty-1 (AT5784)
PRJEB37252 Col-0 (AT6909)

## Footnotes

1 Regions with shared ancestry as a result of a speciation event.

2 Regions with shared ancestry as a result of a duplication event.

3 Haplotype is a collection of highly correlated structural variants.

## References

[1] Multiple alignment format. https://genome.ucsc.edu/FAQ/FAQformat.html#format5.

[2] Sebastian Deorowicz, Agnieszka Danek, and Marcin Niemiec. Gdc 2: Compression of large collections of genomes. Scientific reports, 5:11565, 2015.

[3] P. Deutsch. Gzip file format specification version 4.3, 1996.

[4] Alexander Dilthey, Charles Cox, Zamin Iqbal, Matthew R Nelson, and Gil McVean. Improved genome inference in the mhc using a population reference graph. Nature genetics, 47(6):682–688, 2015.

[5] John Ellson, Emden Gansner, Lefteris Koutsofios, Stephen C North, and Gordon Woodhull. Graphviz—open source graph drawing tools. In International Symposium on Graph Drawing, pages 483–484. Springer, 2001.

[6] Brendan J Frey and Delbert Dueck. Clustering by passing messages between data points. science, 315(5814):972–976, 2007.

[7] Markus Hsi-Yang Fritz, Rasko Leinonen, Guy Cochrane, and Ewan Birney. Efficient storage of high throughput dna sequencing data using reference-based compression. Genome research, 21(5):734–740, 2011.

[8] Erik Garrison, Jouni Sirén, Adam M Novak, Glenn Hickey, Jordan M Eizenga, Eric T Dawson, William Jones, Shilpa Garg, Charles Markello, Michael F Lin, et al. Variation graph toolkit improves read mapping by representing genetic variation in the reference. Nature biotechnology, 2018.

[9] Faraz Hach, Ibrahim Numanagić, Can Alkan, and S Cenk Sahinalp. Scalce: boosting sequence compression algorithms using locally consistent encoding. Bioinformatics, 28(23):3051–3057, 2012.

[10] Lin Huang, Victoria Popic, and Serafim Batzoglou. Short read alignment with populations of genomes. Bioinformatics, 29(13):i361–i370, 2013.

[11] Zamin Iqbal, Mario Caccamo, Isaac Turner, Paul Flicek, and Gil McVean. De novo assembly and genotyping of variants using colored de bruijn graphs. Nature genetics, 44(2):226, 2012.

[12] Daniel C Jones, Walter L Ruzzo, Xinxia Peng, and Michael G Katze. Compression of next-generation sequencing reads aided by highly efficient de novo assembly. Nucleic acids research, 40(22):e171–e171, 2012.

[13] Szymon M Kielwbasa, Raymond Wan, Kengo Sato, Paul Horton, and Martin C Frith. Adaptive seeds tame genomic sequence comparison. Genome research, 21(3):487–493, 2011.

[14] Davis E. King. Dlib-ml: A machine learning toolkit. Journal of Machine Learning Research, 10:1755–1758, 2009.

[15] Heng Li. A proposal of the graphical fragment assembly format. http://lh3.github.io/2014/07/19/a-proposal-of-the-grapical\\-fragment-assembly-format, 2014.

[16] Jasper Linthorst, Marc Hulsman, Henne Holstege, and Marcel Reinders. Scalable multi whole-genome alignment using recursive exact matching. bioRxiv, 2015.

[17] David J Lipman and William R Pearson. Rapid and sensitive protein similarity searches. Science, 227(4693):1435–1441, 1985.

[18] Bo Liu, Hongzhe Guo, Michael Brudno, and Yadong Wang. debga: read alignment with de bruijn graph-based seed and extension. Bioinformatics, 32(21):3224–3232, 2016.

[19] Sorina Maciuca, Carlos del Ojo Elias, Gil McVean, and Zamin Iqbal. A natural encoding of genetic variation in a burrows-wheeler transform to enable mapping and genome inference. In International Workshop on Algorithms in Bioinformatics, pages 222–233. Springer, 2016.

[20] Shoshana Marcus, Hayan Lee, and Michael C Schatz. Splitmem: a graphical algorithm for pan-genome analysis with suffix skips. Bioinformatics, 30(24):3476–3483, 2014.

[21] Eugene W Myers. The fragment assembly string graph. Bioinformatics, 21(suppl 2):ii79–ii85, 2005.

[22] Ngan Nguyen, Glenn Hickey, Daniel R Zerbino, Brian Raney, Dent Earl, Joel Armstrong, W James Kent, David Haussler, and Benedict Paten. Building a pan-genome reference for a population. Journal of Computational Biology, 22(5):387–401, 2015.

[23] Adam M Novak, Erik Garrison, and Benedict Paten. A graph extension of the positional burrows-wheeler transform and its applications. In International Workshop on Algorithms in Bioinformatics, pages 246–256. Springer, 2016.

[24] Adam M Novak, Glenn Hickey, Erik Garrison, Sean Blum, Abram Connelly, Alexander Dilthey, Jordan Eizenga, MA Saleh Elmohamed, Sally Guthrie, André Kahles, et al. Genome graphs. bioRxiv, page 101378, 2017.

[25] Idoia Ochoa, Mikel Hernaez, and Tsachy Weissman. idocomp: a compression scheme for assembled genomes. Bioinformatics, 31(5):626–633, 2014.

[26] Benedict Paten, Mark Diekhans, Dent Earl, John St John, Jian Ma, Bernard Suh, and David Haussler. Cactus graphs for genome comparisons. Journal of Computational Biology, 18(3):469–481, 2011.

[27] Benedict Paten, Adam M Novak, Jordan M Eizenga, and Erik Garrison. Genome graphs and the evolution of genome inference. Genome research, pages gr–214155, 2017.

[28] Benedict Paten, Adam M Novak, Erik Garrison, and Glenn Hickey. Superbubbles, ultrabubbles and cacti. In International Conference on Research in Computational Molecular Biology, pages 173–189. Springer, 2017.

[29] Armando J Pinho and Diogo Pratas. Mfcompress: a compression tool for fasta and multi-fasta data. Bioinformatics, 30(1):117–118, 2013.

[30] Armando J Pinho, Diogo Pratas, and Sara P Garcia. Green: a tool for efficient compression of genome resequencing data. Nucleic acids research, 40(4):e27–e27, 2011.

[31] Benjamin Raphael, Degui Zhi, Haixu Tang, and Pavel Pevzner. A novel method for multiple alignment of sequences with repeated and shuffled elements. Genome Research, 14(11):2336–2346, 2004.

[32] Korbinian Schneeberger, Jörg Hagmann, Stephan Ossowski, Norman Warthmann, Sandra Gesing, Oliver Kohlbacher, and Detlef Weigel. Simultaneous alignment of short reads against multiple genomes. Genome biology, 10(9):R98, 2009.

[33] Jeremy John Selva and Xin Chen. Srcomp: Short read sequence compression using burstsort and elias omega coding. PloS one, 8(12):e81414, 2013.

[34] Claude E Shannon and Warren Weaver. The mathematical theory of communication. 1948.

[35] Jared T Simpson and Richard Durbin. Efficient construction of an assembly string graph using the fm-index. Bioinformatics, 26(12):i367–i373, 2010.

[36] Congmao Wang and Dabing Zhang. A novel compression tool for efficient storage of genome resequencing data. Nucleic Acids Research, 39(7):e45, 2011.

[37] Daniel R Zerbino and Ewan Birney. Velvet: algorithms for de novo short read assembly using de bruijn graphs. Genome research, 18(5):821–829, 2008.

[38] Yongpeng Zhang, Linsen Li, Yanli Yang, Xiao Yang, Shan He, and Zexuan Zhu. Light-weight reference-based compression of fastq data. BMC bioinformatics, 16(1):188, 2015.

[39] Yu Zhang and Michael S Waterman. An eulerian path approach to local multiple alignment for dna sequences. Proceedings of the National Academy of Sciences of the United States of America, 102(5):1285–1290, 2005.

